# Organic availability and microbial competition for acetate suppress methane emissions during the conversion of gypsum in sewage sludge

**DOI:** 10.64898/2026.06.20.733556

**Authors:** Gage R. Coon, Vieyiti Kouadio, Cameron W. M. Murphy, Hui Sun, Oliver Jagoutz, Tanja Bosak

## Abstract

Conventional anaerobic digestion emits methane from organic waste. Here, we investigate a sulfate-based alternative that suppresses methane production and generates alkaline solutions that may sequester carbon by carbonate precipitation. Although methanogenesis is known to occur when reduced organic carbon is replete and sulfate is limiting, it remains unclear whether methane emissions during microbial conversion of waste gypsum are primarily driven by community composition or organic availability. By comparing fluxes of electrons from organic matter toward sulfate or methane in microbial communities grown on different organic loads, we show that community composition, microbial growth, and organic availability collectively determine sulfide and methane fluxes. Lower organic loads increase the importance of syntrophic interactions with fermenters and competition between sulfate reducing bacteria and methanogens due to scarcity of substrates. Microbes present in the original sewage sludge reduce less sulfate, produce more methane, and generate less alkalinity compared to the communities enriched by multiple cycles of growth in the presence of sulfate and sewage sludge. The inoculation of communities enriched at low organic loadings in the presence of sulfate decreases the production of methane by enabling the growth of sulfate reducing bacteria from the order *Desulfobacterales* that can oxidize acetate to CO_2_ and compete with methanogens for acetate. The use of such enrichments in sludge treatment systems can stimulate the removal of organic substrates and waste gypsum, while suppressing methane production, over timescales comparable to those in the current sludge treatment systems that do not contain sulfate.

## Introduction

To meet global phosphorus demands for fertilizer production, phosphate rock (apatite) is treated with sulfuric acid to produce phosphoric acid. This process generates phosphogypsum, a waste product that contains primarily gypsum (CaSO_4_ × 2H_2_O) with trace impurities such as fluoride and radioactive elements such as uranium and thorium. The storage and treatment of phosphogypsum present environmental challenges: approximately 300 million tons are produced annually, and 86% is either stacked indefinitely or discharged into the ocean^1^. Flue gas desulfurization and titanium dioxide production generate additional waste gypsum^2^. The associated disposal issues have motivated the search for economically viable strategies to valorize waste gypsum.

Proposed solutions—such as incorporating waste gypsum into road materials or using it as a soil amendment—face environmental and scalability challenges^3^. An alternative approach is gypsum bioconversion that involves microbial sulfate reduction (MSR), in which sulfate-reducing bacteria (SRB) convert sulfate from gypsum to sulfide, releasing calcium and alkalinity into solution^4^. MSR oxidizes molecular hydrogen or simple organic molecules such as acetate or lactate to reduce sulfate, as shown in the reaction:

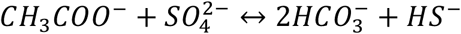

The resulting calcium- and sulfide-rich solution can then be used in downstream processes: sulfide can be converted to elemental sulfur by abiotic or biological oxidation^5–11^, and calcium and alkalinity can precipitate as carbonate minerals, offering a route for carbon sequestration.

A major challenge in implementing this process is the availability of low-cost electron donors for MSR^4,8,12^. Municipal sewage presents a promising solution^4^. In modern wastewater treatment, sewage sludge is typically digested anaerobically, using microbes to convert organic matter into methane via methanogenesis. This methane is often vented or flared to CO_2_, contributing to greenhouse gas emissions^13^.

Extensive research has focused on optimizing microbial communities for anaerobic digestion of sewage sludge through methanogenesis due to its widespread use in wastewater treatment plants; key operational controls include inoculum-to-substrate ratios, upward fluid velocity to retain biomass, and substrate recirculation ratios^14–20^. By adding gypsum directly to anaerobic digesters, electron donors could be redirected from methane production to sulfate reduction. Integration of waste gypsum bioconversion and sulfate reduction into existing sewage treatment infrastructure may help mitigate methane emissions and generate alkaline solutions^21–26^, but requires studies of anaerobic digestion of sewage sludge through MSR.

Prior work demonstrated bioconversion of waste gypsum using sewage sludge, but did not characterize the microbial community that mediated this process^4^. Other research has optimized carbon loading conditions to influence the balance between methanogenesis and sulfate reduction^27–31^, though largely without characterizing the community structure. A recent study has quantified bioconversion kinetics in an optimized reactor using nutrient-rich medium instead of sewage sludge^8^. Other efforts have characterized microbial communities and inoculum source effects in synthetic nutrient-rich systems^32,33^. In general, inoculum compositions and enrichment processes are either unreported or not connected to functional outcomes. Most research to date also does not measure methane production or characterize the microbial community during gypsum treatment on organic sources other than sewage sludge^8,12,32–42^ or reports^43–46^ methane production on sewage sludge without attempting to suppress or relate it systematically to the rates of sulfate reduction, alkalinity generation, or calcium concentrations in the solution.

The rates of organic compound oxidation and subsequent methane and sulfide production are driven by the abundance of organic substrates, collectively referred to as organic loading. This loading is a well-established control on anaerobic redox processes: sulfate reducing bacteria typically outcompete methanogens for electron donors under conditions of low organic loading^47^, where input is often very limited between nearly 0 and 1.5 g COD/L. In contrast, environments with high organic loading—such as eutrophic sediments or sulfate-rich sewage sludge with up to 90 g COD/L—can support both metabolisms simultaneously^48–51^. However, how electron donors are partitioned between these competing pathways in systems with high organic loading and how to control this partitioning remains unclear.

Here, we address these gaps by identifying a microbial community that rapidly reduces sulfate using organic electron donors from municipal wastewater, generates alkalinity, and minimizes methane production. This is done by comparing the evolution of solution chemistry at high and low organic loading in systems inoculated with two distinct microbial enrichments and systems that contain microbes from the original sewage sludge as the only starting community. Analyses of microbial diversity reveal a relationship among the taxa present in the cultures, the removal of organic matter, and the observed rates of sulfate reduction and methane production. The addition of enriched inocula at specific organic loading to sulfate ratios can be used to control microbial competition and scale up the inherently unsterile sludge treatment systems. These insights inform the design of a gypsum bioconversion process that mitigates methane emissions by favoring sulfate reduction and maximizing alkalinity generation.

## Materials and Methods

### Sewage media

Thickened sewage sludge and final treated effluent (FEFF) were sourced from Deer Island Wastewater Treatment Plant, operated by the Massachusetts Water Resources Authority (MWRA). Thickened sewage sludge had a chemical oxygen demand (COD) ranging from 74 to 92 g/L. A concentrated sewage medium, referred to as “high organic loading” conditions, was prepared anaerobically by a 1:1 dilution of sewage sludge with FEFF and sieved through a 0.35 mm mesh. A dilute sewage medium, referred to as “low organic loading” conditions, was prepared anaerobically by another 30.7x dilution of the concentrated sewage medium, resulting in a final 61.4x dilution. The media with high and low organic loading were amended by 22.6 g/L and 5.18 g/L of pure gypsum, respectively. These concentrations are equivalent to a total sulfate concentration of 131.3 mM and 30.1 mM, respectively, resulting in molar COD : sulfate ratios of ∼45 and ∼1.5 at high and low organic loading, respectively. The headspace in the vials was anaerobic (98% N_2_ and ∼2% H_2_)

### Enrichments

Two communities were previously enriched on sewage media in the presence of sulfate, one in the presence of low and one in the presence of high organic loading. These communities had different initial environmental inoculums.

Enriched Community #1 was established by adding sulfide-rich sediment from Plum Island, MA, USA, to sewage medium with 2.5 g COD/L amended with 20 mM sodium sulfate (molar COD: sulfate ≈ 3.9). Sulfide and sulfate concentrations were monitored during each generation of enrichment. Cultures were transferred at 10% v/v to the freshly prepared medium with the same organic and sulfate loading when sulfide and sulfate concentrations began to plateau. Enriched Community #1 used in this manuscript was harvested after three transfers.

Enriched Community #2 was inoculated by a sample from a sulfide-rich stream from a sulfur mine in Sicily, Italy. This sample was added to ∼260x diluted (molar COD: sulfate ≈ 0.4) and filtered sewage with flue-gas desulfurization gypsum. Sulfide concentrations were monitored during the first generations of enrichment, and a new generation was inoculated on ∼260x diluted and 0.8 µm filtered sewage once sulfide concentrations began to plateau. This enrichment process was repeated ten times and as experiments continued^52^.

### Inoculations and incubations

Enrichment cultures of microbes were prepared for inoculation into sewage medium by incubation in the same media as those used in subsequent experiments for two weeks. All incubations were conducted in triplicate with 10% v/v inoculum and incubated at 32°C, shaking at 135 rotations per minute (rpm) in the dark. The initial biomass in the inoculated and uninoculated sewage media could not be determined due to the presence of native microbial communities and organic particulates in the sewage. To monitor the growth of these pre-cultures, sulfide concentrations were measured regularly until they reached at least 1 mM in any given culture, after which the pre-cultures were inoculated into sewage media at 10% v/v. Sulfide concentrations after the inoculation of inocula were 0.16–0.29 mM at high organic loading and 0.47–0.75 mM at low organic loading. Triplicate cultures that contained only sewage media with different organic loadings were not inoculated by any pre-cultures to enable the assessment of importance of different inoculums and organic loadings. The results of the high organic loading experiment are reported up to 8 days of incubation, and results of the low organic loading experiment are reported up to 17 days of incubation.

### Inhibiting sulfate reduction

To test whether acetate was an intermediate in sewage-digesting anaerobic cultures, we quantified its concentration in batch cultures inoculated by homogenized sludge from a bioreactor that received sludge and sulfate at 2.8 COD: sulfate ratio. The sludge was inoculated at 10% v/v into triplicate serum bottles that contained 40 mL of thickened sewage sludge diluted by tap water in a 1:20 ratio and amended by 2.58 g/L of pure gypsum, yielding 6.27 g COD/L after inoculation and a final molar COD: sulfate ratio of ∼13. All incubations were conducted for 16 days at 30°C in the dark. Control triplicate cultures were amended by 5 mM sodium 2-bromoethanesulfonate (BES) to inhibit methanogenesis or 3 mM sodium molybdate to inhibit sulfate reduction. The headspace in all bottles was 32 mL.

### Sulfide concentrations

Sulfide concentrations were measured by the Cline assay^53^ described at dx.doi.org/10.17504/protocols.io.6qpvrq1oolmk/v1. Generally, samples were diluted 100x in 0.05M zinc acetate and measured immediately.

### Methane concentrations

Methane concentrations were measured by a Shimadzu GC-2014 gas chromatograph with a PDHID and helium carrier gas. The pressure in the culture vials with high organic loading was relieved by occasional degassing. We presume any increase in pressure during that time frame is negligible and thus assume 1 atm. Dissolved methane ([CH_4aq_], in mM) was calculated using a mass-balance equation to account for both the headspace and residual aqueous methane at equilibrium:

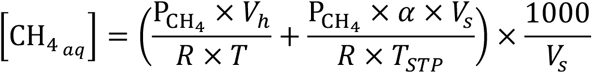

where P_CH4_ is the partial pressure of methane (atm, converted from ppm assuming 1 atm total pressure), *V_h_* is the headspace volume (L), *R* is the universal gas constant (L×atm/mol×K), *T* is the temperature (K), α is the Bunsen solubility coefficient for methane at the experimental temperature (α = 0.029;^54^), T_STP_ is standard temperature (273.15 K), *V_s_* is the volume of the medium containing sewage and gypsum (L), and 1000 is the conversion factor for mM.

### Alkalinity and pH

Alkalinity was measured using a ThermoScientific Orion Total Alkalinity Test Kit (Cat. No. 700010TS). The pH was measured using Vernier Tris-compatible flat pH sensors or AtlasScientific EZO pH sensors.

### Ion concentrations

Sulfate, calcium, and magnesium were measured using a dual-channel Shimadzu ion chromatograph. Cations were measured using the Shodex “Monovalent and Divalent Cation Standards (4) (YK-421) (Phosphoric Acid Eluent)” method, and anions using the Shodex “Anion Standards (19) Rapid Analysis (SI-90 4E)” method, with the column temperature set to 40°C.

### Calculating electron partitioning towards sulfate reduction and methanogenesis

The partitioning of reducing equivalents towards sulfate reduction was monitored by calculating *f* as the molar ratio of sulfide produced over the sum of sulfide produced and methane produced. The partitioning of reducing equivalents towards methanogenesis, the other process of terminal oxidation, as 1 minus *f*, where *f* is the previously calculated ratio towards sulfate reduction. For high-organic loading conditions, *f* was calculated using the measurements of sulfide and methane between the third and the eighth day of incubation (n = 2 – 3 replicates). Relative percent increase in the partitioning of reducing equivalents towards sulfate reduction over this partitioning in the cultures that were not inoculated by any enrichments is the quotient of the respective *f* values towards sulfate reduction.

### Total and volatile solids (TS and VS)

TS and VS were measured using Standard Methods 2540 SOLIDS (23^rd^ ed, 2017). The sludge extracted after the incubation of the cultures was dried at 105°C. Total solids ratio was calculated as the dry mass of the sample normalized to the total wet mass of the sample. VS powder was prepared by anaerobically drying the entire incubation slurry at the end of the experiment. VS was combusted at 550°C and the resulting mass loss was normalized to the dry mass. Liquid was not separated from solids before drying. “% VS removal” is calculated as (mass of initial VS - mass of final VS)/mass of initial VS.

### Calcium to total S ratio (Ca: Total S)

The ratio of calcium to total sulfur was calculated by dividing measured calcium concentrations by the sum of sulfate and sulfide concentrations. To track potential carbonate precipitation, we used the Ca: Total S ratio at day 6 as a reference point and scaled it by changes in total sulfur to calculate an expected ratio over time. Measured Ca: Total S values were then normalized to this expectation, with values below 1 interpreted as indicative of calcium removal, potentially due to carbonate precipitation. Calcium carbonate is the thermodynamically favored precipitate under our conditions, and no magnesium carbonates were detected by XRD (Figure S2), so Ca: Total S was used to assess carbonate mineral formation. Ratios were only calculated from day 6 onward after gypsum fully dissolved and expected stoichiometric Ca: SO_4_^2-^ ratios were observed.

### 16S rRNA gene sequencing

DNA was extracted using the DNeasy PowerSoil Pro Kit (Qiagen, Hilden, Germany) and quantified with Qubit dsDNA HS Assay Kit and Qubit 3 fluorometer (Thermo Fisher Scientific, Waltham, MA, USA). Sequencing was performed on an Illumina MiSeq sequencer using universal primers U515F (5’-GTGCCAGCMGCCGCGGTAA-3’) and E786R (5’-GGACTACHVGGGTWTCTAAT-3’), targeting the V4 region. Sequences were processed with the DADA2 pipeline, and taxonomy was assigned using the MiDAS reference database version 5.3. Raw sequence data are available at ENA project accession PRJEB89716, and the full analysis code is available at https://github.com/gagecoon/OM_loading_PG.

## Results

We hypothesized that concentrated sewage sludge with added gypsum would support both sulfate reduction and methanogenesis, but that the activities of these competing metabolisms would depend on the composition of microbial communities that are influenced by the addition of preexisting inoculums. We further hypothesized that the addition of external inoculums would be less consequential at high organic loading, when abundant organic substrates could supply reducing equivalents to both sulfate reduction and methanogenesis. To test these hypotheses, we incubated triplicate cultures that tested three different conditions. Control cultures contained only concentrated sewage sludge (∼45 g COD/L) amended with excess gypsum. The untreated sewage sludge contained the original microbial community that had had no prior exposure to sulfate-rich conditions. The same sludge and microbes were present in separate triplicate cultures that contained the same sewage medium, but these cultures were additionally inoculated by Enrichment Culture #1 or #2 that contained sulfate-reducing bacteria enriched by serial passaging on sewage in the presence of sulfate (see Methods).

All cultures simultaneously reduced sulfate and produced methane in the presence of high organic loading. Both communities that were inoculated by previous enrichments reduced more sulfate in 8 days than the original community (Table 1), suggesting a shift in the microbial function to favor gypsum conversion in the presence of high organic loading. Enriched communities also generated 3–15% more alkalinity than the original community in the sewage (Figure S1B).

**Table 1:** Methanogenesis and sulfate reduction under high organic loading conditions.

| Enrichment | sulfate<br>reduced<br>(mM) | methane<br>produced<br>(mM) | % electrons towards |  | increase in sulfate<br>reduction vs<br>original |
| --- | --- | --- | --- | --- | --- |
|  |  |  | sulfate<br>reduction | methanogenesis |  |
| #1 | 5.5 ± 1 | 3.4 ± 0.7 | 62 ± 9 | 38 ± 9 | +38% |
| #2 | 3 ± 1.2 | 2.2 ± 0.4 | 56 ± 10 | 44 ± 10 | +24% |
| Original | 2.4 ± 0.2 | 3.0 ± 1.0 | 45 ± 6 | 55 ± 6 |  |

To reduce methane production and divert reducing equivalents toward sulfate reduction, we lowered the organic loading and hypothesized that the suppression of methanogenesis would depend on community composition of the inoculum. To test this, we incubated triplicate cultures of the two enriched communities and the original community in diluted sewage sludge (molar COD : sulfate of ∼1.5) with added gypsum. This design allowed us to assess the influences of microbial community composition in both enrichments and sewage on sulfate reduction and methane production in solutions with low organic loading. Both enriched communities reduced sulfate more rapidly than the original community (Fig. 1) and generated more alkalinity. Enriched Community #1 and Enriched Community #2 reduced sulfate at similar initial rates, but Enriched Community #2 reduced sulfate faster after the first three days of incubation (Figs. 1A). Dilution of the sewage sludge reduced methane production in all cultures (Fig. 1B), but only Enriched Community #2 did not produce any methane (Fig. 1B). Thus, enrichment conditions can influence the composition of the microbial communities that, when inoculated onto appropriately diluted sewage sludge in the presence of sulfate, can reduce sulfate while not producing methane (Fig. 2).

**Figure 1:**
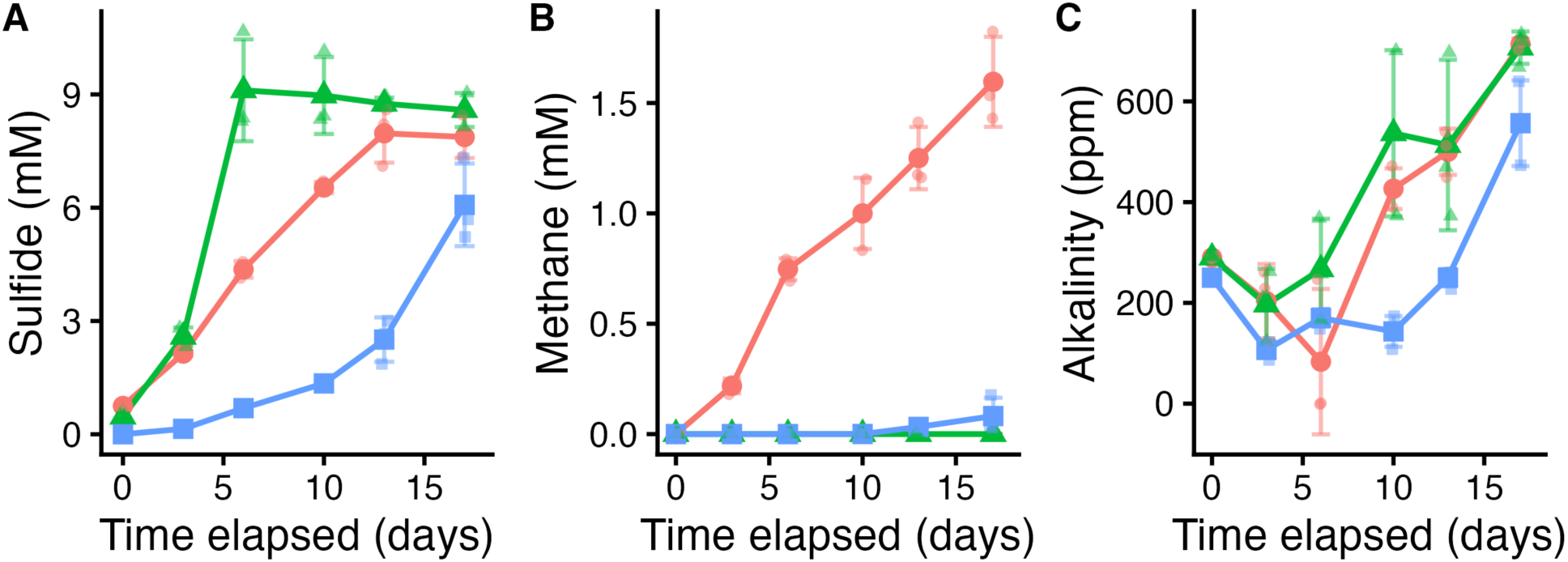
Evolution of solution chemistry in enriched and original sewage communities. A) average sulfide concentrations, B) average methane concentrations, C) average alkalinity concentrations (ppm as CaCO_3_). Red circles: Enriched Community #1, green triangles: Enriched Community #2, blue squares: original community in sewage. Large symbols show average values from triplicate serum bottles, the error bars are standard deviation. Small lighter symbols show raw values from each replicate from the three conditions marked by red (Enriched Community#1), green (Enriched Community #2) and blue (original community in sewage).

**Figure 2:**
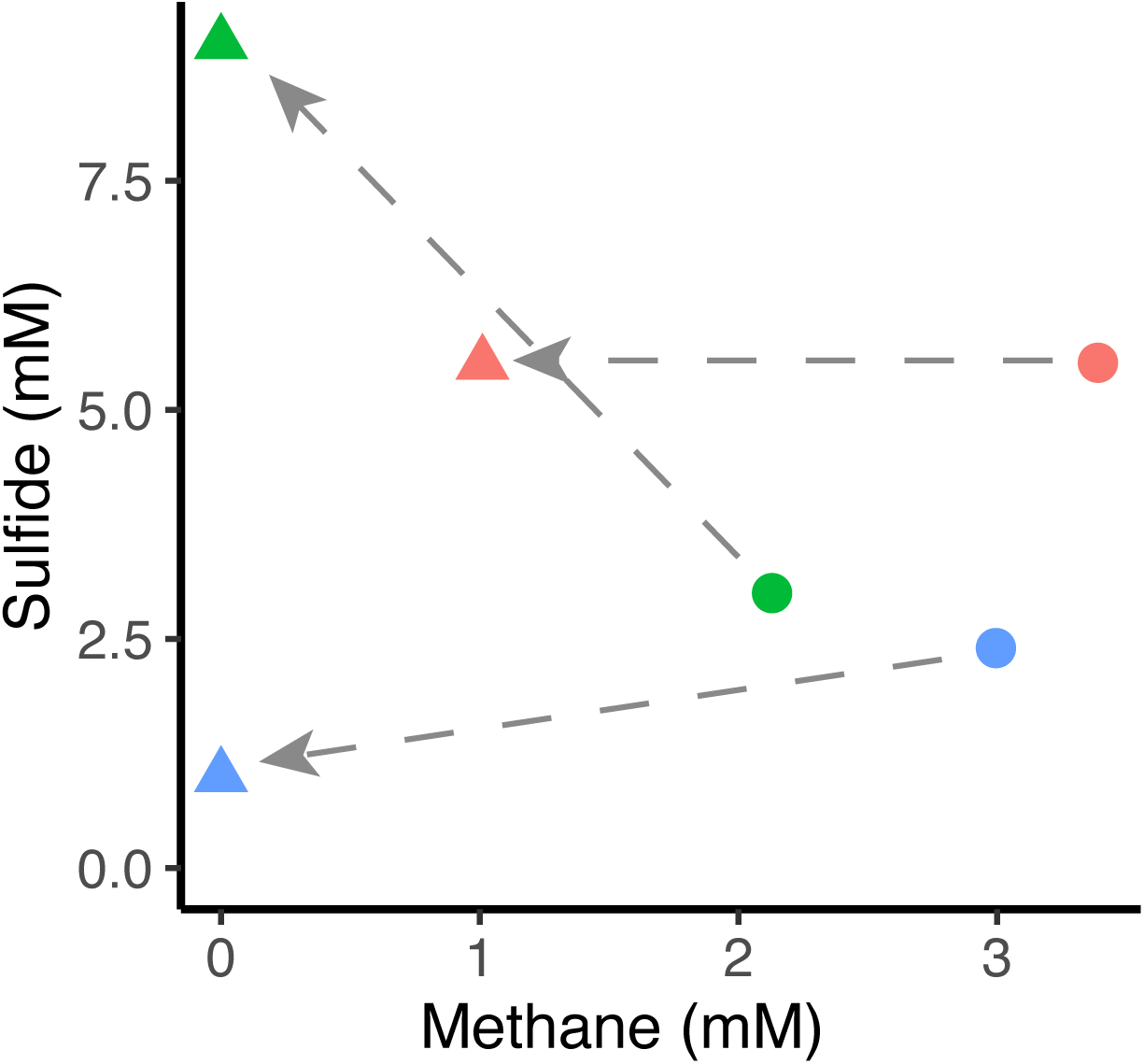
Effect of sewage dilution from molar COD : sulfate ∼45 to ∼1.5 on average methane and sulfide concentrations. Triangles show cultures grown on diluted and circles show cultures grown on undiluted sewage sludge. Red symbols show Enriched Community #1, green symbols show Enriched Community #2, and blue symbols show the original community. All data points show average values from triplicate culture bottles after eight days of incubation.

Efficient removal of organic matter remains a central goal of wastewater treatment. In addition to measuring sulfide, methane, and alkalinity concentrations, we assessed the degradation of organic matter by measuring the mass of volatile solids (VS) and quantified the concentration of calcium and alkalinity in the solution. VS removal is a key performance metric in wastewater treatment because it reflects the extent of organic degradation and assesses the performance of the sludge treatment process^55^. We quantified VS after the cessation of sulfate reduction in the cultures with low organic loading (molar COD: sulfate of ∼1.5). Both enriched communities removed more VS than the original community (Table 2). Assuming COD reduction is proportional to VS removal, this suggests an estimated COD removal of ∼36% for Enriched Community #1 and ∼29% for Enriched Community #2, roughly doubling the ∼14% removal observed in the original sewage community. This is consistent with enhanced microbial activity due to the addition of pre-enriched inoculums. These results suggest that communities enriched to degrade sewage sludge (Fig. 1) degraded sludge more efficiently and released more alkalinity.

**Table 2:** Volatile solid (VS) removal at low organic loading.

| Enrichment | % VS removal | % VS removal<br>versus original |
| --- | --- | --- |
| Enriched<br>#1 | 36.3 ± 3.2 | +146.9% |
| Enriched<br>#2 | 29.0 ± 10.2 | +97.3% |
| Original | 14.7 ± 6.7 |  |

Microbes generated less alkalinity in the presence of low organic loading (Fig. 1C) than in medium with high organic loading (Fig. S1), while releasing more calcium into the solution (Fig. 3B, Supplemental Data). Because calcium is released only when sulfate is reduced, carbonate formation under experimental conditions described here depends primarily on sulfate reduction.

**Figure 3:**
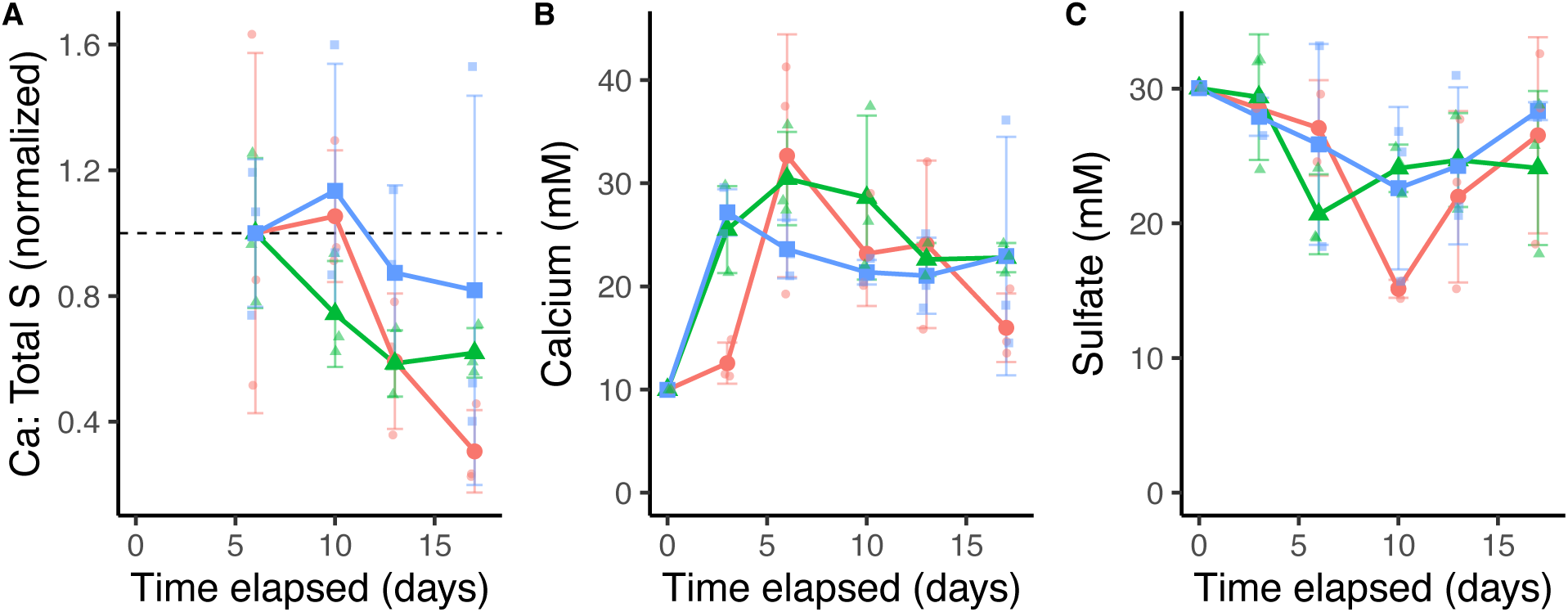
Comparison of calcium and sulfate concentrations at molar COD: sulfate of ∼1.5. A) Average ratio of calcium to total sulfur (Ca: Total S) normalized to expected ratio (dashed line) determined by the release of equimolar concentrations of calcium and sulfate due to the dissolution of gypsum. Values below the dashed line show a calcium deficit. B) Average calcium concentration. C) Average sulfate concentration. Red circles show Enriched Community #1, green triangles show Enriched Community #2, blue squares show the original sewage community. All large symbols show average values measured in triplicate serum bottles, the error bars are standard deviation. Small, lighter symbols with the same colors show raw values from each respective replicate culture bottle.

Next, we wanted to identify organic intermediates that can be used by both methanogens and sulfate reducers. To do so, we inoculated nine serum bottles with sewage sludge and gypsum (Methods) with a mature inoculum from a continuous sulfate-reducing digester amended with sewage sludge and gypsum at molar COD: sulfate of ∼13 so sulfate was limiting. Three bottles were incubated as experimental controls, three were amended by 5 mM BES to inhibit methanogenesis and three were amended by 3 mM molybdate to inhibit sulfate reduction (see Methods). Because acetate has been identified as the main organic intermediate between MSR and methanogenesis in organic-rich systems^56,57^, and because we did not detect any hydrogen when measuring methane concentrations by gas chromatography, we hypothesized that this small organic acid would also accumulate in the bottles where either methanogenesis or MSR were inhibited. The initial medium contained negligible concentrations of acetate (µM), so all concentrations of acetate measured over time show how much acetate accumulated in inhibited and uninhibited cultures.

As expected, both MSR and methanogenesis could operate without inhibition in control cultures and acetate concentrations in those cultures were higher at all times than the concentrations in the cultures where sulfate reduction was inhibited. Communities without inhibitors reduced the supplied ∼20 mM sulfate entirely in eight days (Fig. 4A). The rates of methane production in these cultures increased 2.5-fold after all sulfate was reduced (Fig. 4), suggesting competition between MSR and methanogenesis and demonstrating the limitation of MSR by sulfate rather than by organic electron donors. Inhibiting sulfate reduction by adding molybdate increased the production of methane by more than an order of magnitude, once again suggesting competition between SRB and methanogens for electron donors (Fig. 4B). The concentration of acetate was lower in the cultures where sulfate reduction was inhibited (Fig. 4D), showing the potential of methanogens to consume acetate. The inhibition of methanogenesis also lowered acetate concentrations compared to the control cultures at five days of incubation (Fig. 4D). Sulfate reduction removed all sulfate after eight days from the cultures where methanogenesis was inhibited by BES (Fig. 4A) and the concentration of acetate increased (Fig. 4D), likely due to the continuing production of acetate by anaerobic fermenting microbes.

**Figure 4:**
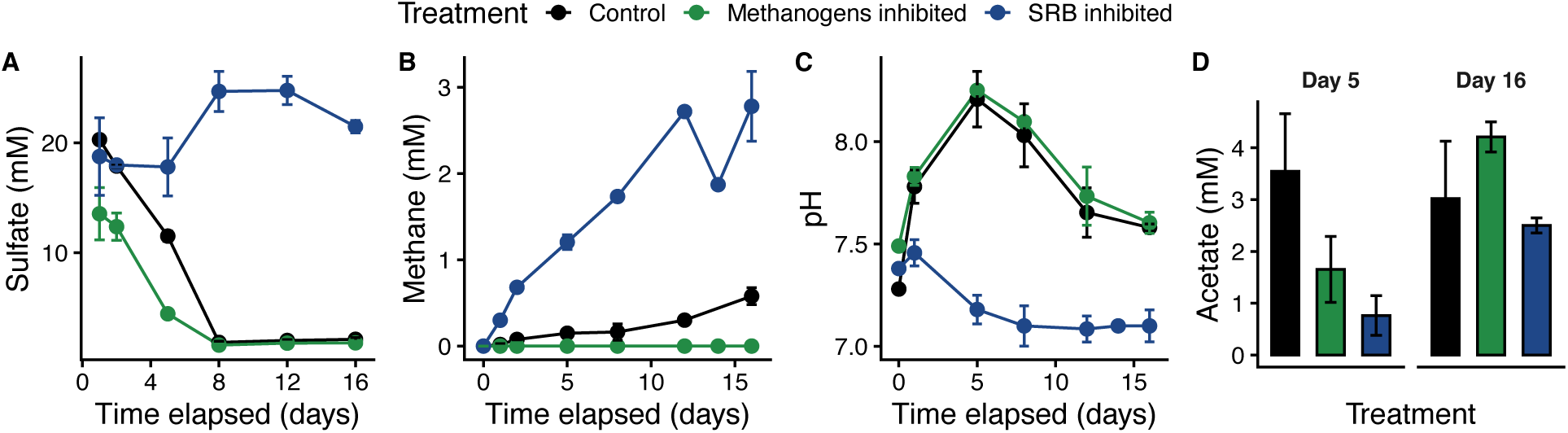
Effects of sulfate reduction and methanogenesis on solution chemistry in batch cultures incubated on sewage medium with a molar COD: sulfate ratio of ∼13. Concentrations of A) sulfate (mM), B) methane (mM), C) pH, and D) acetate (mM) throughout the 16-day incubation. Error bars represent the standard deviation for triplicate cultures.

To determine the metabolism responsible for the changes in pH and alkalinity, we measured the solution pH throughout the experiment and found nearly identical values in sulfate-reducing cultures (BES-inhibited cultures) and controls (Fig. 4C). Therefore, MSR increased the pH, but methanogenic activity did not.

In the attempt to relate the differences in sulfate reduction, methane production, and the removal of volatile solids to the structure of microbial communities, we sequenced the 16S rRNA gene of all enrichment cultures and the original community grown at a molar COD: sulfate ratio of ∼1.5 (Figs. 5, 6). This allowed us to compare the compositions of microbial communities as a function of experimental conditions, with a focus on the microbial groups involved in fermentation, sulfate reduction, and methanogenesis. We compared the communities sampled after 17 days of incubation. At that time, sulfate reduction ceased in the cultures amended by microbial enrichments (Fig. 1), but one of the enriched communities still produced methane (Fig. 1). These data enable comparisons of microbial communities in the presence of low organic loading.

**Figure 5:**
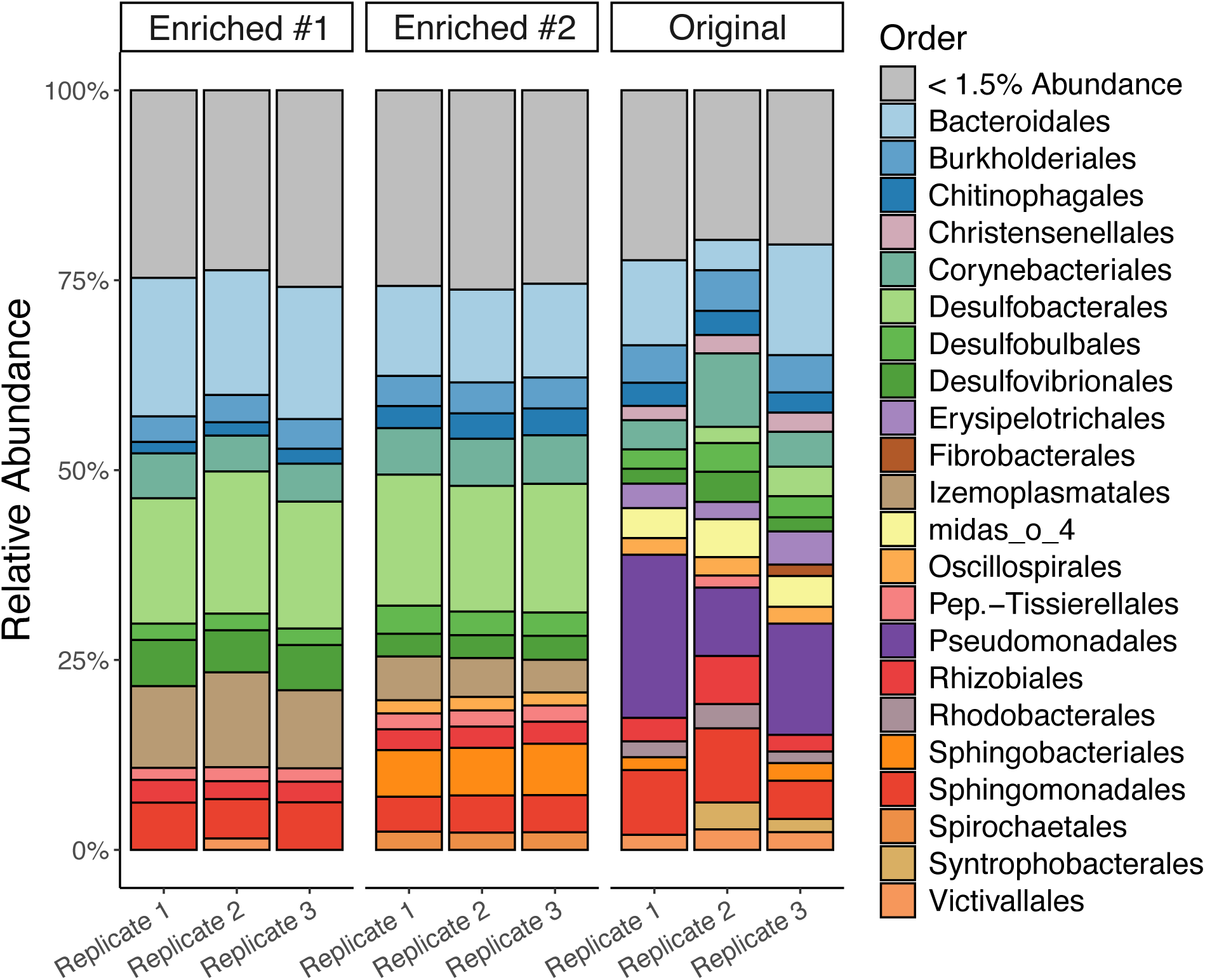
Orders represented by the 16S rRNA sequences from triplicate cultures of enriched and original sewage communities.

**Figure 6:**
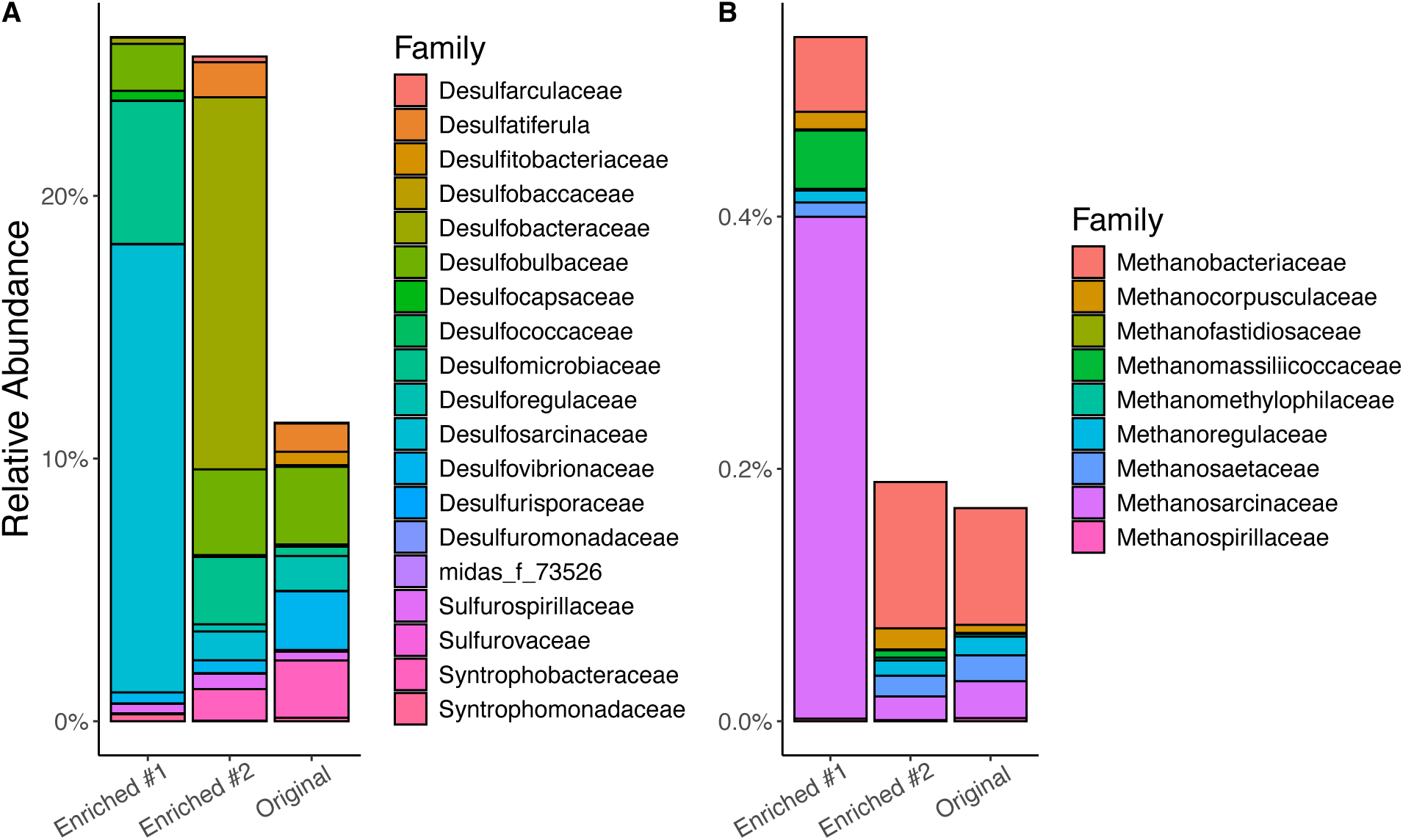
Average compositions of microbial families represented by the 16S rRNA sequences from triplicate cultures of enriched and original communities. A) Sulfate-reducing bacteria, B) Methanogenic archaea.

In spite of different enrichment conditions and starting inoculums, both enriched communities converged toward a similar taxonomic composition at the order level (Fig. 5). Community profiles were consistent across biological triplicates, indicating that selective pressures shaped reproducible community structures even in the complex, variable, non-sterile background of sewage sludge. The abundances of different families changed as a function of enrichment procedures relative to the native sewage community. *Chitinophagales* was present at similar levels across all conditions, but the two enriched communities contained more *Desulfobacterales* than the original community (Figs. 5, 6). *Pseudomonadales* was highly abundant in the original community but nearly absent from both enriched communities, *Synergistales* and *Sphingomonadales* were also more abundant in the original community. Several known fermentative orders were abundant in all conditions, including *Bacteroidales*, *Christensenellales* and *Peptostreptococcales–Tissierellales*. *Bacteroidales* were the most abundant fermenting microbes in both enriched communities. *Izemoplasmatales* and *Desulfovibrionales* were also much more abundant in both enriched communities relative to the original community (Fig. 5). In contrast, *Syntrophobacterales*, *Syntrophibacterales*, *Eubacterales*, *Flavobacterales*, *Fibrobacterales*, and *Pseudomonadales* were less abundant in Enriched Community #1 compared to the original community. Enriched Community #2 contained lower abundances of *Fibrobacterales*, *midas_o_4*, *Pseudomonadales* relative to the original community, and higher abundances of *Sphingobacteriales* relative to Enriched Community #1 and the native sewage community. These results suggest that the enrichment procedure changed the fermentative community and the abundances of sulfate reducing bacteria while preserving a core community structure dominated by *Bacteroidales*.

To further relate these taxonomic shifts to the observed rates of sulfate reduction and methanogenesis, we analyzed the relative abundances of sulfate reducing and methanogenic families (Fig. 6). Enriched Community #1 contained the 16S rRNA sequences of sulfate reducers and methanogens. The latter sequences in this community were about three times more abundant than in Enriched Community #2 or the original community. In contrast, Enriched Community #2, contained more sequences of sulfate reducing bacteria, but not methanogens, than the original community. *Desulfomicrobiaceae* were present in both enriched communities, but more abundant in Enriched Community #2. Enriched Community #1 was also enriched in *Desulfosarcinaceae* and contained fewer sequences of *Desulfitobacteriaceae*, *Desulfobulbaceae*, *Desulfatiferula* and *Desulforegulaceae*. Sequences of *Syntrophobacteraceae* were nearly absent from Enriched Community #1 but present in both Enriched Community #2 and the original community. Enriched Community #2 was further enriched in *Desulfobacteraceae*. Several other sulfate reducing families, including *Desulfurisporaeceae*, were detected only in trace amounts across all conditions.

Methanogens exhibited low abundance across all treatments, but their community compositions changed with experimental conditions. The relative abundance of methanogens was the highest in Enriched Community #1 (∼0.6%) and comprised mostly of *Methanosarcinaceae* and *Methanomassiliicoccaceae*. Enriched Community #2 and the original community instead contained primarily *Methanobacteriaceae* (∼0.2%). Enriched Community #2 contained somewhat more *Methanobacteriaceae* and fewer *Methanosarcinaceae* relative to the original community. These results suggest that fermentation and oxidation of organic matter in Enriched Community #2 provided electrons to sulfate, but not to an abundant or metabolically versatile community of methanogens. These findings show that enrichment strategies can reproducibly shift microbial communities in sewage sludge and promote sulfate reduction over methanogenesis in the presence of dilute sewage sludge and waste gypsum.

## Discussion

### Enrichments and decreased organic loading divert more electron donors to sulfate reduction

Sulfate reducing microbial communities can effectively convert gypsum to sulfide and alkalinity using dilute municipal sewage sludge as the source of electron donors for MSR. The addition of enrichment cultures and the dilution of concentrated sewage sludge can stimulate organic removal by microbial reduction of sulfate from gypsum with negligible methane emissions. Organic removal rates were enhanced by the addition of enrichments. The methane suppression observed here is consistent with the values reported by other studies using synthetic media and compost^58–62^. but had not been quantified previously in gypsum-converting systems using municipal sewage sludge. Diverting reducing equivalents toward sulfate reduction requires balancing microbial competition and regulating organic loading and COD to sulfate ratios. Although higher organic loadings (molar COD: sulfate ≈ 45) allow gypsum bioconversion, they also provide electron donors to methanogens, resulting in comparable rates of methane generation and sulfate reduction. Lower organic loading (molar COD: sulfate ≈ 1.5) mitigates this competition and reduces methane emissions in appropriately enriched and conditioned microbial systems.

### Syntrophy and competition for substrates

The availability of organic substrates and syntrophic interactions between fermenters and terminal oxidizers shape the composition and function of microbial communities enriched in the presence of sewage sludge and sulfate. Fermenters are the most abundant organisms in the anaerobic communities (Figure 5) and constitute the metabolic foundation that supports terminal metabolisms through the production of volatile fatty acids (VFAs), hydrogen and alcohols^63^. Dominant fermentative orders observed here, *Bacteroidales*, *Christensenellales*, and *Peptostreptococcales–Tissierellales*, are common in anaerobic digesters^64^ and likely mediate the primary degradation of proteins and carbohydrates into acetate, propionate, butyrate, and lactate^65–68^. Increased abundances of *Bacteroidales* and *Izemoplasmatales* in both enriched communities indicate enhanced hydrolysis and acidogenesis^69^, suggesting that the structure of the fermentative community fundamentally constrains terminal electron accepting pathways. Typically, VFAs do not accumulate significantly in stable anaerobic digesters^70^, suggesting the close coupling of fermentation kinetics and subsequent terminal metabolisms that oxidize organic products of fermentation.

Distinct taxonomic profiles of sulfate reducers and methanogens across communities reflect differences in substrate availability and metabolic preferences. In Enriched Community #1, the continuous production of methane during MSR coincides with increased abundance of metabolically versatile methanogens, particularly *Methanosarcinaceae* and *Methanomassiliicoccaceae*. *Methanosarcinaceae* can utilize acetate, hydrogen, and methanol^71^, while *Methanomassiliicoccaceae* are obligate methylotrophic methanogens reliant on hydrogen-dependent reduction of methanol^72^. The increased abundances of these taxa in Enriched Community #1 suggest that the availability of diverse fermentation byproducts favors methanogens capable of exploiting multiple substrates. In contrast, the most abundant methanogens in Enriched Community #2 and the original unenriched sewage community belong to the hydrogenotrophic family *Methanobacteriaceae*, which generates methane by oxidizing H_2_^73,74^. The negligible methane production in these communities likely reflects limitation by H_2_ and competitive exclusion by sulfate reducers.

The decline in secondary fermenters—e.g., acetogens that convert the less competitive non-acetate VFAs to acetate and H_2_, such as *Syntrophobacterales*—in Enriched Community #1 suggests rapid consumption of VFAs by terminal reducers, reducing the necessity for syntrophic oxidation of intermediates such as propionate to acetate. The competition for electron donors by SRB and methanogens under our experimental conditions relies on acetate, suggesting that the differences in methane suppression between Enriched Community #1 and 2 stem from the varying affinities of SRB for acetate. The dominance of the sulfate reducing family *Desulfosarcinaceae* in Enriched Community #1 provides a possible mechanistic explanation: these sulfate reducers preferentially oxidize substrates such as propionate and certain aromatic compounds that cannot be used by methanogens. This expected lack of competition for organic substrates may explain the observed coexistence of SRB that cannot oxidize acetate and acetoclastic methanogens such as *Methanosarcinaceae*. Some species of *Desulfosarcinaceae* can oxidize acetate, but they are considered a metabolically versatile group that oxidize a wider variety of organic compounds compared to the canonical acetate-oxidizing *Desulfobacteraceae* abundant in Enriched Community #2. Additionally, *Methanomassiliicoccaceae*, an obligate hydrogen-dependent methylotrophic methanogen^72^, are more abundant in Enriched Community #1. This trend likely reflects the availability of hydrogen and methanol provided by fermenters and minimal competition of *Methanomassiliicoccaceae* and *Desulfosarcinaceae* for hydrogen. Although *Desulfosarcinaceae* can engage in syntrophy with anaerobic methane-oxidizing archaea (ANME)^75^, their co-occurrence here with abundant methanogens is likely due to complementary, non-competitive substrate use. In contrast, Enriched Community #2 contains primarily the acetate-oxidizing *Desulfobacteraceae* that directly compete with methanogens for this substrate. Thus, early establishment of substrate-competitive SRB in Enriched Community #2 may suppress methanogenic activity, shifting the microbial community to favor the reduction of sulfate. These sulfate reducers are prevalent in the initial inoculum. These results highlight the importance of competition for substrates among fermenters, methanogens, and sulfate reducers in the shaping of communities that mediate gypsum conversion using electron donors from sewage (Fig. 7).

**Figure 7:**
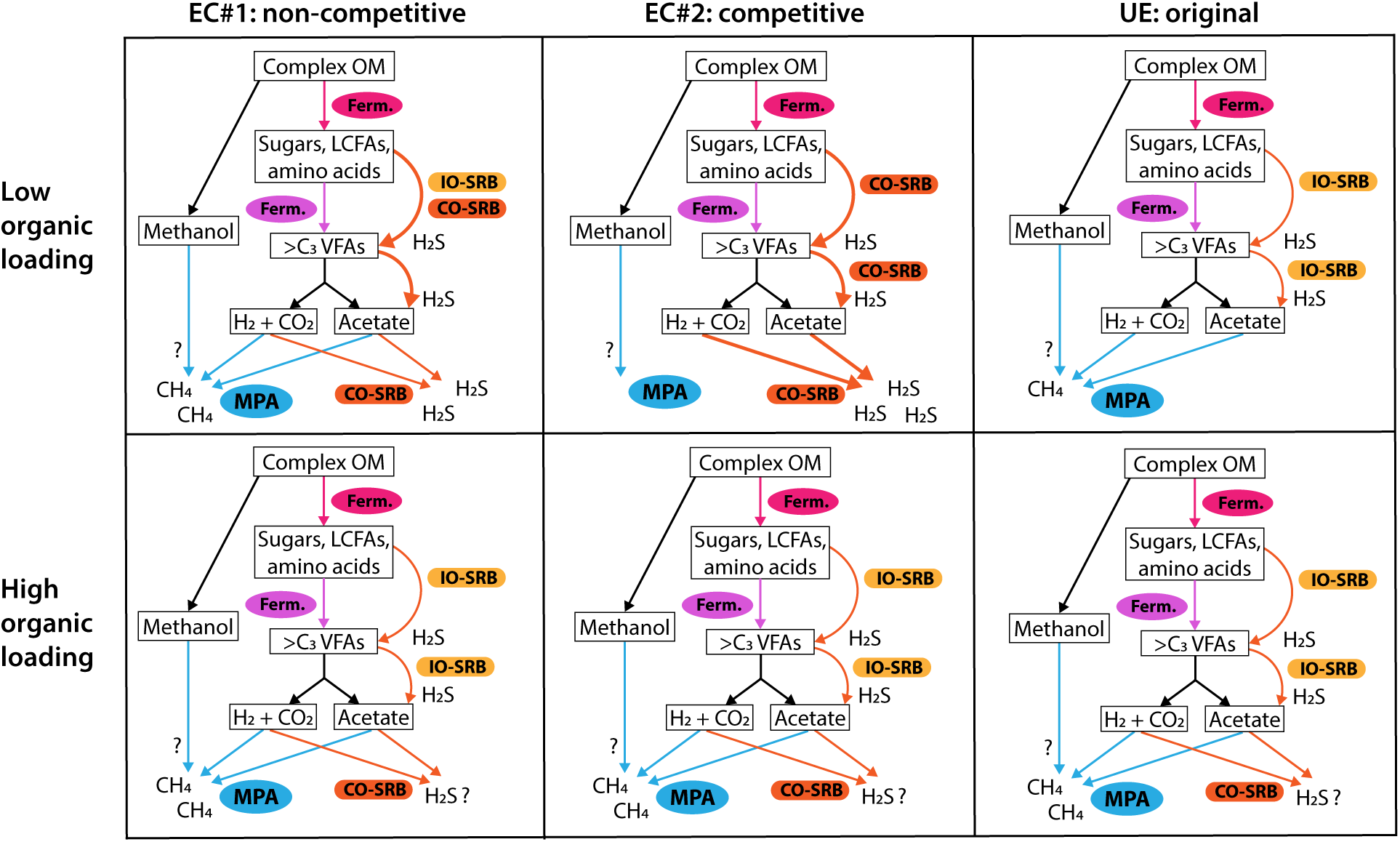
Schematic of the inferred microbial interactions that govern the partitioning of electrons. Low organic loading ≈ 1.5 COD: sulfate (molar ratio), high organic loading ≈ 45 COD: sulfate (molar ratio). EC#1 = inoculated with enriched community #1, EC#2= inoculated with enriched community #2, UE = native sewage community that encounters sulfate for the first time. Colored arrows denote specific microbial activities: pink (fermentation), blue (methanogenesis by methane-producing archaea, MPA), yellow (incomplete sulfate reduction by IO-SRB), and orange (complete sulfate reduction by CO-SRB). Abbreviations: OM, organic matter; VFAs, volatile fatty acids; >C3, organic compounds with carbon chains longer than three; Ferm, fermentative microorganisms; LCFAs, long-chain fatty acids.

Together, our experiments and observations support a model wherein both thermodynamic considerations and competitive interactions among microbes that reduce sulfate or produce methane control the balance between sulfate reduction and methanogenesis. The latter interactions depend on the competition for substrates produced by anaerobic fermenting microbes. In the system described here, acetate emerges as the main such substrate. Microbial metabolic competition is increasingly more important at lower organic loading due to the general scarcity of organic substrates. Thus, lowering the organic loading by dilution provides a lever with which to suppress methane production in anaerobic digesters. Enrichments for robust sulfate reduction and organic removal rates at lower organic loadings can reduce the production of this potent greenhouse gas during the bioconversion of sewage sludge and waste gypsum.

### Scalability and Community Transferability

Directing electron donors toward sulfate reduction without significant methane production could leverage existing wastewater treatment infrastructure. Nearly all dominant taxa in the enriched communities were also detected in the original community present in sewage sludge, but the successive cycles of cultivation on gypsum and sewage enhanced their abundances. So the addition of external diversity during the enrichment process is likely not necessary, and enrichment processes could occur in anaerobic digesters and then continually transferred. Enriched communities removed more volatile solids (VS), but it remains unclear if VS removal will be sufficient in large sulfate-reducing digesters with semicontinuous operation. The community that produced methane removed more VS in the same time frame than the community that produced little methane (Table 2), so the desired VS removal in bioreactors may require a balance between partial methanogenesis and sulfate reduction. Optimized flow-through systems may enhance VS removal and should be the focus of future work. While large dilutions of livestock manure are economically unfavorable for gypsum conversion^31^, municipal sewage may be a better option because it is often thickened 25-50 times before digestion. Avoiding this step, and instead adding gypsum earlier in the treatment process, could achieve the ∼60x dilution explored herein. Alternatively, seawater presents a viable post-thickening diluent, providing additional sulfate and cations without significant disruption to existing wastewater infrastructure. A similar approach has been implemented in Hong Kong, where the SANI (Sulfate reduction, Autotrophic denitrification and Nitrification Integrated) process effectively treats sulfate-rich wastewater^21–26^. Similar approaches can be adopted for gypsum conversion and alkalinity generation.

Enrichment of sulfate-reducing communities at low organic loading reduced sulfate at rates equivalent to ∼250 mg gypsum/L/day while suppressing methane, driven by the enrichment of *Desulfobacterales*. Future work should evaluate whether similar outcomes can be achieved through in-situ enrichment in continuous reactor systems and whether the extent of organic removal is sufficient under continuous operation.

## Acknowledgements

This project was supported by the Chan-Zuckerberg and Siegelman funds to the Advanced Carbon Mineralization Initiative (ACMI). GRC thanks fellowship funding from GE Vernova. TB acknowledges funding by MIT School of Science Fundamental Science Investigator award. We extend acknowledgement to Lisa Wong and colleagues at Deer Island’s Wastewater Treatment Plant for sample collection. This work was carried out in part through the use of MIT.nano’s facilities.

