## Supplemental Materials for "Organic availability and microbial competition for acetate suppress methane emissions during the conversion of gypsum in sewage sludge"

**Supplementary data**

High organic loading experiments generated sulfide and alkalinity and plateau after day 8 of incubation (Figure S1).

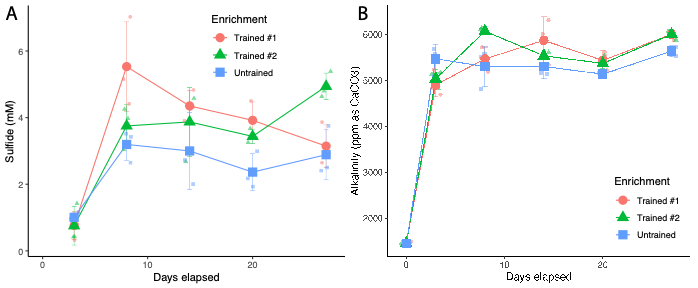

Figure S1: Solution chemistry of triplicate incubations of trained and untrained communities under high organic loading conditions. (A) Average sulfide concentrations used to estimate sulfate reduction rates. (B) Average methane concentrations. Error bars represent standard deviation.

Magnesium carbonate precipitates were not detected via XRD of solids dried (where precipitation of carbonates is strictly abiotic, Figure S2).

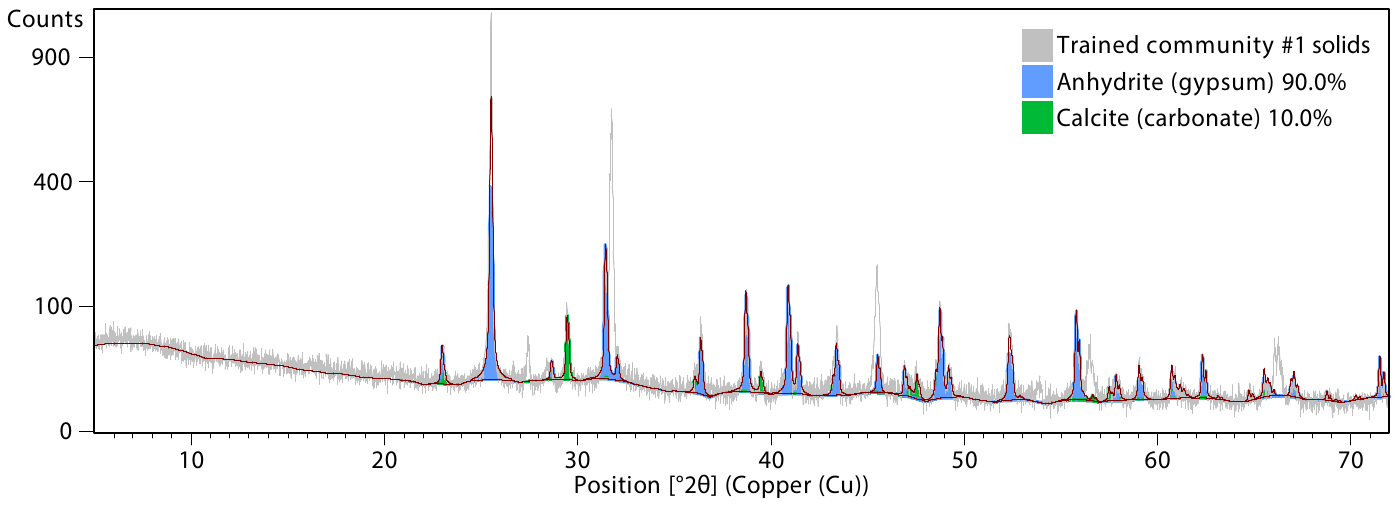

Figure S2: Representative XRD of precipitated solids.
